# Narrow-sense heritability estimation of complex traits using identity-by-descent information

**DOI:** 10.1101/164848

**Authors:** Luke M. Evans, Rasool Tahmasbi, Matthew Jones, Scott I. Vrieze, Gonçalo R. Abecasis, Sayantan Das, Doug W. Bjelland, Teresa R. deCandia, - Haplotype Reference Consortium, Gonçalo Abecasis, David Altshuler, Carl A Anderson, Andrea Angius, Jeffrey C Barrett, Sonja Berndt, Michael Boehnke, Dorrett Boomsma, Kari Branham, Gerome Breen, Chad M Brummett, Fabio Busonero, Harry Campbell, Peter Campbell, Andrew Chan, Sai Chen, Emily Chew, Massimiliano Cocca, Francis S Collins, Laura J Corbin, Francesco Cucca, Petr Danecek, Sayantan Das, Paul I W de Bakker, George Dedoussis, Annelot Dekker, Olivier Delaneau, Marcus Dorr, Richard Durbin, Aliki-Eleni Farmaki, Luigi Ferrucci, Lukas Forer, Ross M Fraser, Timothy Frayling, Christian Fuchsberger, Stacey Gabriel, Ilaria Gandin, Paolo Gasparini, Christopher E Gillies, Arthur Gilly, Leif Groop, Tabitha Harrison, Andrew Hattersley, Oddgeir L Holmen, Kristian Hveem, William Iacono, Amit Joshi, Hyun Min Kang, Hamed Khalili, Charles Kooperberg, Seppo Koskinen, Matthias Kretzler, Warren Kretzschmar, Alan Kwong, James C Lee, Shawn Levy, Yang Luo, Anubha Mahajan, Jonathan Marchini, Steven McCarroll, Mark I McCarthy, Shane McCarthy, Matt McGue, Melvin McInnis, Thomas Meitinger, David Melzer, Massimo Mezzavilla, Josine L Min, Karen L Mohlke, Richard M Myers, Matthias Nauck, Deborah Nickerson, Aarno Palotie, Carlos Pato, Michele Pato, Ulrike Peters, Nicola Pirastu, Wouter Van Rheenen, J Brent Richards, Samuli Ripatti, Cinzia Sala, Veikko Salomaa, Matthew G Sampson, David Schlessinger, Robert E Schoen, Sebastian Schoenherr, Laura J Scott, Kevin Sharp, Carlo Sidore, P Eline Slagboom, Kerrin Small, George Davey Smith, Nicole Soranzo, Timothy Spector, Dwight Stambolian, Anand Swaroop, Morris A Swertz, Alexander Teumer, Nicholas Timpson, Daniela Toniolo, Michela Traglia, Marcus Tuke, Jaakko Tuomilehto, Leonard H Van den Berg, Cornelia M van Duijn, Jan Veldink, John B Vincent, Uwe Volker, Scott Vrieze, Klaudia Walter, Cisca Wijmenga, Cristen Willer, James F Wilson, Andrew R Wood, Eleftheria Zeggini, He Zhang, Jian Yang, Michael E. Goddard, Peter M. Visscher, Matthew C. Keller

## Abstract

Heritability is a fundamental parameter in genetics. Traditional estimates based on family or twin studies can be biased due to shared environmental or non-additive genetic variance. Alternatively, those based on genotyped or imputed variants typically underestimate narrow-sense heritability contributed by rare or otherwise poorly-tagged causal variants. Identical-by-descent (IBD) segments of the genome share all variants between pairs of chromosomes except new mutations that have arisen since the last common ancestor. Therefore, relating phenotypic similarity to degree of IBD sharing among classically unrelated individuals is an appealing approach to estimating the near full additive genetic variance while avoiding biases that can occur when modeling close relatives. We applied an IBD-based approach (GREML-IBD) to estimate heritability in unrelated individuals using phenotypic simulation with thousands of whole genome sequences across a range of stratification, polygenicity levels, and the minor allele frequencies of causal variants (CVs). IBD-based heritability estimates were unbiased when using unrelated individuals, even for traits with extremely rare CVs, but stratification led to strong biases in IBD-based heritability estimates with poor precision. We used data on two traits in ~120,000 people from the UK Biobank to demonstrate that, depending on the trait and possible confounding environmental effects, GREML-IBD can be applied successfully to very large genetic datasets to infer the contribution of very rare variants lost using other methods. However, we observed apparent biases in this real data that were not predicted from our simulation, suggesting that more work may be required to understand factors that influence IBD-based estimates.

## INTRODUCTION

The proportion of phenotypic variance due to additive genetic variation, termed narrow-sense heritability (*h*^2^), is perhaps the most fundamental aspect of a trait’s genetic architecture and has both medical and evolutionary significance (Visscher *et al.*, 2008;; Tenesa and Haley, 2013). Traditionally, *h*^2^ has been estimated from family-based studies 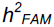 which have suggested that for many complex traits, much of the phenotypic variance is due to additive genetic variance (Polderman *et al.*, 2015). However, 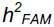 estimates may be biased by factors shared by close relatives, such as non-additive genetic and common environmental effects (Eaves *et al.*, 1978; Coventry and Keller, 2005; Yang *et al.*, 2010; Zuk *et al.*, 2012; Tenesa and Haley, 2013). Alternatively, the variance explained by genetic markers in unrelated individuals, for instance from genome-wide association studies (GWAS), may avoid many of the possible confounding factors of family-based studies. However, the thousands of variants associated with complex traits (Visscher *et al.*, 2012 Ripke *et al.*, 2014 Wood *et al.*, 2014) often explain only a fraction of trait heritability, with the difference often called the “missing heritability.” This missing heritability may result from causal variants (CVs) that are rare or otherwise poorly tagged by commercial genotyping arrays, insufficient sample sizes to detect small effect variants that do not reach genome-wide significance, and biases that inflate 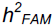.

Recently, methods have been developed to estimate the phenotypic variance explained by all genotyped markers simultaneously in unrelated individuals, 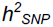 (Yang *et al.*, 2010; Speed *et al.*, 2012; Bulik-Sullivan *et al.*, 2015). Most of these approaches generally use a genetic relatedness matrix (GRM) that reflects allele sharing or the average correlation between individuals *i* and *j* across genotyped SNPs with entries:

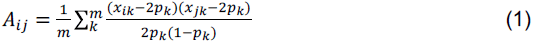

where *m* is the number of SNPs, *x*_*jk*_ is the genotype (coded as 0, 1, or 2) of individual *j* at the *k*^*th*^ locus, and *p*_*k*_ is the minor allele frequency (MAF) of the *k*^*th*^ locus. The variance-covariance of the phenotype is

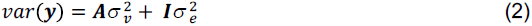

where the variance explained by the SNPs 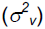 and error variance 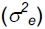 are estimated using restricted maximum likelihood (REML). The method, termed GREML, is implemented in packages such as GCTA (Yang, *et al.*, 2011). We refer to matrix **A** (of dimension *n* x *n* and with elements *A*_*ij*_) as the “SNP-GRM.” The proportion of the variance explained by all SNPs is an estimate of “SNP-based heritability” 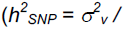 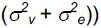. By using unrelated individuals, these approaches avoid the confounding of non-additive genetic and environmental effects that can occur in family or twin-based studies, and by estimating all marker effects jointly, the contribution from variants with small effect sizes is captured. Using marker-based approaches, 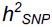 estimates for some complex traits, such as height, have approached 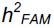 suggesting that little of the heritability remains missing (Yang *et al.*, 2015). For other traits, such as BMI, schizophrenia and neuroticism, 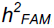 estimates remain larger than 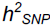 and a substantial amount of the heritability remains “still missing” (Lee *et al.*, 2012; Yang *et al.*, 2015).

Advances on the original approach by Yang *et al.* (2010) have better captured the effects of rare CVs and account for linkage disequilibrium (LD) of markers across the genome, leading to increased 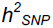 estimates (Yang *et al.*, 2015). However, even with the best-performing methods such as MAF- and LD-stratified GREML (GREML-LDMS) and large imputation reference panels, downward bias is likely. Imputation quality declines at low MAF, resulting in a downward bias when causal variants are very rare (MAF<0.0025) and for diverse populations underrepresented in sequencing panels (Evans *et al.*, 2017). The underestimation of variance due to rare CVs may partly explain why 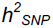 remains below 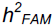 for many traits. Thus, developing alternative methods to estimate the variation caused by very rare variants is an important goal.

One such alternative is to leverage information on the proportion of the genome shared identical-by-descent (IBD) between pairs of individuals in a sample (Visscher *et al.*, 2006;; Hayes *et al.*, 2009; Zuk *et al.*, 2012; Browning and Browning, 2013), and use a GRM whose elements are the estimated proportions of IBD between all pairs of individuals (IBD-GRM) to estimate heritability 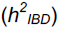 This is in some ways similar to classical family-based estimates of heritability, which are based on the expected proportion of the genome shared IBD between close relatives (Falconer and Mackay, 1996; Lynch and Walsh, 1998; Visscher *et al.*, 2006). However, rather than using close relatives, an appealing alternative is to estimate pairwise IBD segments directly between all pairs of unrelated (or technically, distantly related) individuals in a sample and use these estimated relationship values to estimate the additive genetic variation. Such an IBD-based approach should capture additive genetic variation due to all but the rarest variants and, so long as close relatives have been removed from the sample, the IBD-based *h*^2^ estimate should be uncontaminated by confounding factors shared by close relatives. Note that we use “IBD”to denote two homologous chromosomal segments that came from the same common ancestor without intervening recombination, such that the sequence identity of the two segments is identical except at sites where new mutations arose since the last common ancestor. The probability that such mutations arose is a function of the time since the last common ancestor, and therefore a function of the length of the shared IBD segment (Wakeley, 2009). Very long segments are therefore more likely to be identical at all sequence sites, whereas shorter ones are more likely to have occasional differences at sites harboring typically very rare variants. Thus, IBD-GRMs calculated from increasingly long IBD thresholds should capture sharing at increasingly rare CVs.

Such IBD-based GRMs have been used in several instances to estimate heritability. Price *et al.* (2011) and Zaitlen *et al.* (2013) used IBD segments in an Icelandic dataset with close relatives to estimate heritability in quantitative and disease traits, leveraging the known familial relationships within the Icelandic cohort to identify IBD segments. While they demonstrated that IBD could be used for heritability estimation, using close relatives leads to possible confounding of shared environmental or non-additive genetic effects, as noted above. Indeed, Zaitlen *et al.* (2013) found higher heritability estimates using closer relatives, consistent with confounding from non-additive genetic and/or shared environment effects. Using simulated data, Zuk *et al.* (2012) demonstrated that the slope estimated from regressing phenotypic similarity (defined as the standardized phenotypic product of individuals *i* and *j*, Z_i_ x Z_j_) on the IBD-GRM elements from long IBD segments—known as Haseman-Elston (H-E) regression—provides an unbiased estimate of the additive genetic variance in isolated founder populations. Browning and Browning (2013) estimated IBD tracts in a Finnish cohort of 5,400 individuals, and used the resulting IBD GRM in both H-E regression and GREML to estimate 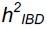 for nine quantitative metabolic traits. 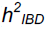 was higher than 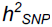 for only five of the nine traits, and never significantly so. The most notable result of their study was the over two-fold higher standard errors for 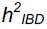 (~0.17) compared to 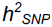 (~0.07), due to the lower variation in the off-diagonal elements of the IBD-GRM compared to the SNP-GRM, suggesting that very large sample sizes will be required to obtain meaningful results in non-founder populations.

Several important questions about IBD-based heritability estimation remain in light of these findings. First, can an IBD-based approach account for very rare CVs? Previous studies (e.g., Browning and Browning, 2013) have simulated CVs from SNPs present on genotyping arrays, which are more common, have generally higher LD than most variants throughout the genome, and are shared across ancestry groups, and therefore do not provide an accurate picture of how *h*^2^ estimation methods perform when CVs do not share these same properties. Thus, it is unclear whether 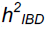 estimates are unbiased estimates of *h*^2^ in the presence of rare CVs. Second, the studies mentioned above utilized isolated founder populations that were both more homogeneous and more related than non-founder populations. To what extent does genetic stratification bias 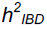, and how feasible are such IBD-based method in samples from non-founder populations, which are much more readily available?

To address these questions, we used thousands of recently sequenced whole genomes from the Haplotype Reference Consortium (McCarthy *et al.*, 2016) to simulate phenotypes under a range of conditions, including various genetic architectures and levels of stratification, then estimated narrow-sense heritability (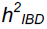) using the an IBD-GRM, either alone or in combination with various SNP-based GRMs. By simulating CVs from whole genome sequences rather than commercial array SNPs, our study was able to examine the role of all but the rarest frequency classes of CVs in the genome under realistic genomic conditions. We then estimated 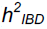 for height and BMI in the UK Biobank with over 120,000 individuals.

## MATERIALS AND METHODS

### Samples and Population Structure

We tested the 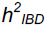 estimation method using simulated phenotypes derived from Haplotype Reference Consortium (HRC) whole genome sequence data (McCarthy *et al.*, 2016). Full details of the HRC can be found in McCarthy *et al.* (2016). Briefly, this resource comprises roughly 32,500 individual whole genome sequences from multiple sequencing studies, with phased genotypes with a minor allele count of at least 5 at all sites. This large sequence dataset allowed us to simulate CVs across all MAF classes down to ~.0003 with real patterns of LD (within and among chromosomes). It also allowed us to simulate SNP markers available on existing commercial genotyping arrays in order to mimic the process of IBD detection in SNP data. We obtained permission to access the following HRC cohorts (recruitment region & sample size): AMD (Europe & worldwide; 3,189), BIPOLAR (European ancestry; 2,487), GECCO (European ancestry; 1,112), GOT2D (Europe, 2,709), HUNT (Norway; 1,023), SARDINIA (Sardinia; 3,445),TWINS (Minnesota; 1,325), 1000 Genomes (worldwide; 2,495), UK10K (UK;; 3,715) (see (McCarthy *et al.*, 2016) for additional details of the HRC). This set of cohorts, which included isolated subpopulations of European descent, allowed investigation into the effects of stratification on estimates. The subset totaled 21,500 whole genome sequences comprising 38,913,048 biallelic SNPs. This is the same set of individuals and simulated phenotypes used in Evans *et al.* (2017) to compare SNP-based heritability methods. Below, we briefly describe our approach.

Our goal was to assess the accuracy and potential bias of the 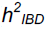 estimation method using data similar to those collected for a typical GWAS analysis and 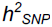 estimation. In order to mimic this kind of data, we first extracted variant positions corresponding to a widely-used commercially available genotyping array, the UK Biobank Affymetrix Axiom array. We then identified individuals of primarily European ancestry, using principal components analysis with 133,603 MAF- and LD-pruned markers (plink2 (Chang *et al.*, 2015) command: --maf 0.05 --indep-pairwise 1000 400 0.2) to identify a grouping associated with the 1000 Genomes European individuals in the HRC. This dataset comprised 19,478 individuals including Finnish and Sardinian samples (Fig. S1).

From within this European ancestry dataset, we identified clusters that contained different levels of genetic heterogeneity within them (Fig. S2). The most structured group contained all samples (N=19,478). The somewhat structured group excluded Sardinian and Finnish samples (N=14,424). The low structure group contained northern/western European samples (N=11,243), and the least structured was a subset of mainly British Isles samples (N=8,506). We used GCTA (Yang, *et al.*, 2011) with LD- and MAF-pruned SNPs to estimate relatedness and remove the minimal number of individuals from pairs with relatedness > 0.1 within each of the four samples. In the most homogeneous and smallest sample with no genetic structure, this left 8,201 individuals.

In order to eliminate the influence of varying sample size in our comparison across the range of stratification, we randomly chose 8,201 of the unrelated individuals from within each of the other three stratification subsamples. We similarly tested a lower relatedness cutoff of 0.05 within each group (leaving 7,792; 8,115; 8,129; and 8,186 individuals for the four subsamples), and used both subsets later to examine how a 0.1 or 0.05 relatedness cutoff influences 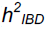 estimates.

### Simulated Phenotypes Using Whole Genome Sequencing Data

To test how 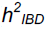 estimation our method performed on a range of genetic architectures, we simulated phenotypes from CVs drawn randomly from five MAF ranges: common (MAF>0.05), uncommon (0.01<MAF<0.05), rare (0.0025<MAF<0.01), very rare (0.0003<MAF<0.0025), and all variants randomly drawn with MAF>0.0003.Phenotypes were generated with 1,000 or 10,000 CVs from the model *y*_*i*_ *= g*_*i*_ *+ e*_*i*_, where *g*_*i*_ = ∑*w* _*ik*_*bb*_*k*_, *w*_*ik*_ is the genotype (coded as 0, 1, or 2) of individual i at the *k*^*th*^ CV, and *bb*_*k*_ is the *k*^*th*^ allelic effect size, drawn from ~N(0,1/[2*p*_*k*_(1-*p*_*k*_)]), where *p*_*k*_ is the MAF of allele *k* within each of the four samples, which assumes larger additive effects for rarer variants. The *g*_*i*_’s were standardized and residual error was added as ~N(0,(1- *h*^2^)/*h*^2^) for a simulated *h*^2^ of 0.5. A total of 400 replications were performed for each CV MAF range and for each of the four stratification subsets.

### Mixed Models for Heritability Estimation

We estimated heritability for each simulation using GCTA (Yang, Lee, *et al.*, 2011). We tested different models to assess our IBD-based GREML method (GREML-IBD). First, we used the single IBD-GRM with GREML to estimate 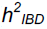 Second, to partition the genetic variance into that tagged by common SNPs and that tagged by haplotype sharing, presumably from rarer CVs, we used a two GRM model (GREML-IBD+SNPs) with the IBD-GRM (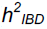) and a common SNP-GRM derived from Axiom array positions with MAF>0.01 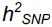. Last, we estimated genetic variances due to LD- and MAF-stratified imputed variant SNP-GRMs (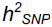) as well as the IBD-GRM (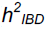) as a comparison to the GREML-LDMS method, which we term GREML-IBD+LDMS. From previous work, we knew that GREML-LDMS underestimates variance attributable to the rarest CVs when using imputed data. We therefore wished to determine if the IBD-GRM could capture that missing heritability. To do this, we estimated 16 SNP-GRMs stratified into the above 4 MAF categories and 4 LD score quartiles using imputed genome-wide variants, and included these plus the IBD GRM in the model (17 GRMs total). To determine if the IBD-GRM captured the genetic variance due to the rarest CVs, we also tested a model with 12 SNP-GRMs, removing the rarest MAF category described above, for a total of 13 GRMs in the analysis (three MAF categories X four LD score quartiles + 1 IBD-GRM). To impute, we first phased SNP data using SHAPEIT2 (Delaneau *et al.*, 2013), imputed using minimac3 (Das *et al.*, 2016), and retained variants with imputation *R*^2^≥0.3 (Yang *et al.*, 2015). We used the HRC sequence data as our imputation reference panel after removing all target (8201 unrelated + relatives) individuals in the HRC reference panel, thereby assuring ~independence (no relatedness) between the target and reference panels. Additional details of the imputation procedure can be found in Evans *et al.* (2017). We estimated LD scores for the LD stratification using GCTA. In all cases we used the –reml-no-constrain option of GCTA, and included 20 principal components (PCs;; 10 from worldwide PC analysis and 10 from the specific subsample PC analysis) as continuous covariates, with sequencing cohort as a categorical covariate.

### Estimating IBD-GRMs

To mimic computationally phased SNP data with realistic phase errors, we first un-phased the sequence data for each data subset and then re-phased the Axiom array positions using SHAPEIT2 (Delaneau *et al.*, 2013). We then used FISHR2 (Bjelland *et al.*, 2017) to identify shared haplotype segments that are putatively IBD across all pairs of individuals within each of our four structure samples. FISHR2 first uses a modified version of GERMLINE (Gusev *et al.*, 2009) to find candidate IBD segments. It then improves the accuracy of the segment endpoints by comparing an observed moving average of haplotype mismatches (potential phase or SNP call errors) for a given candidate IBD segment to (a) the distribution of haplotype mismatches in segments that are almost certainly IBD (the middlemost sections of very long IBD segments) and (b) the distribution of haplotype mismatches in segments that are almost certainly non-IBD (between random pairs of individuals at matched locations). FISHR2 truncates candidate segments when this moving average becomes more consistent with non-IBD than IBD. FISHR2 is more accurate than leading competitors at detecting long (> 3 cM) IBD segments and is the only software that gives unbiased estimates of the true length of IBD segments. The parameters we used for FISHR2 were stringent (command line - err_hom 4 -err_het 1 –min_snp 128 –min_cm_initial 1 –min_cm_final 1 –window 50 – gap 100 -h_extend -w_extend –homoz -emp-ma-threshold 0.06 -emp-pie-threshold 0.015 -count.gap.errors TRUE), chosen to minimize false positive IBD detection (Bjelland *et al.*, 2017). We used an initial length threshold of 1 cM, but because longer IBD segments are more likely to share rare variants, we also identified segments of length greater than 2, 3, 4, 6, 9, and 12 cM. The FISHR2 parameters we used should lead to consistently low false positive rates (<.05) at all threshold lengths, and should lead to a sensitivity that increases as a function of the length of the true IBD segments, and should be >.90 for IBD segments >3cM (Bjelland *et al.*, 2017). To reduce the influence of low recombination regions artificially extending segments (e.g., due to one or a few matching IBS SNPs that are far from the termini of true IBD segments), we windsorized genetic map positions by setting the maximum distance between adjacent markers to 0.2 cM, and used an initial 1 cM minimum IBD segment length threshold.

We then summed the length in Mb of all segments shared between each pair of individuals and divided by twice the length of the genome. This IBD-GRM then represents the estimated proportion of the genome, *D*_*ij*_, shared IBD between individuals *i* and *j* in the sample, similar to the *A*_*ij*_ elements of the SNP-GRM. We created IBD-GRMs for each minimum segment cM length threshold. As recombination rate varies throughout the genome, in two of the subsamples we also tested whether an IBD-GRM based on the summed cM length of segments influences heritability estimates.

### Stratification Effects

We performed four additional analyses to further determine the influence of stratification on 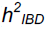 estimates. First, to test whether bias observed in stratified samples was due to inadequate control of structure, we ran K-means clustering on the somewhat stratified subsample for K=2 clusters, then ran PC analysis within each of the two clusters. We included the first 35 PCs within each cluster, for a total of 90 PCs (the original 20 plus 35 from each cluster). Because PC analysis was run within each cluster separately, we set the PC scores for the alternate cluster to 0 (the mean).

Second, we tested, within the stratified subsample, whether including 10 additional PCs from very rare variants could correct for the upward bias (Mathieson and McVean, 2012). We used 150,000 randomly selected very rare SNPs from the WGS data and pruned for LD (plink2 command: --indep-pairwise 1000 400 0.2), leaving 129,710 variants for the PCA. As a comparison, we also estimated heritability with no covariates included.

Third, we estimated 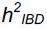 for phenotypes in which all CVs were drawn from odd chromosomes using IBD-GRMs estimated only from the even chromosomes. The presence of uncontrolled cryptic relatedness or population structure can lead to cross-chromosome LD that inflates *h*^2^ estimates (Yang *et al.*, 2011). We estimated the correlation of off-diagonal GRM elements between the IBD-GRMs from even chromosomes and those from odd chromosomes. We also examined the correlation between the off-diagonal elements from IBD-GRMs and the off-diagonal elements from GRMs built from very rare (0.0003<MAF<0.0025) and common (MAF>0.05) sequence variants. This tested whether correlations between even and odd chromosome IBD-GRMs were stronger in more stratified subsamples, and whether the correlation with very rare variants was stronger with increasing minimum cM length of the IBD-GRM.

Last, simultaneously fitting GRMs derived from each chromosome protects against cross chromosome correlations induced by stratification or cryptic relatedness because the estimates of variance explained by one GRM are conditional on the other GRMs (Yang *et al.*, 2011). However, because the variances of the off-diagonal elements in the IBD-GRMs were so small, models with 22 IBD-GRMs would not converge. Instead, we tested a two GRM model with one IBD-GRM estimated from the odd numbered chromosomes and a second from the even numbered chromosomes, which should partially address the effects of long-range LD (Speed *et al.*, 2012).

### Heritability of Complex Traits in the UK Biobank

We applied the IBD-based approaches to height and body mass index (BMI) data in the UK Biobank, a very large resource of ~500K adults from the UK, genotyped using the Affymetrix Axiom array (Sudlow *et al.*, 2015). The current release includes ~150K genotyped individuals, imputed using the combined UK10K/1000 Genomes reference panels. We used this resource previously, and full details on quality control can be found in Evans *et al.* (2017). We identified putative IBD segments as described above using FISHR2 and then calculating IBD-GRMs with minimum cM thresholds of 2, 3, 4, 6, 9, and 12cM. We applied a relatedness cutoff of 0.05, and used individuals of European ancestry, resulting in a final sample size of ~120K individuals included in the analysis (Fig. S2). We used GCTA to estimate variance components and included sex, UK Biobank assessment centre, genotype measurement batch, and qualification (highest level of educational attainment) as categorical covariates, and the Townsend deprivation index, age at assessment, age at assessment squared, and the 15 PC scores from the UK Biobank as quantitative covariates. We compared these models using Akaike information criterion with sample size correction (AICc) (Burnham and Anderson, 2002), and used this to determine if additional information was added by using an IBD-GRM.

## RESULTS

### Simulated Phenotypes – GREML-IBD

Using a single IBD-GRM, 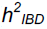 estimates varied greatly depending on the MAF range of the CVs in simulated phenotypes and the amount of stratification in the subsample (Fig. 1). In the two more homogeneous subsamples, 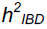 increased then stabilized with increasing IBD segment length threshold. The 95% CI overlapped the true heritability (0.5) for all IBD thresholds > 4 cM and for all CV MAF classes, suggesting that GREML-IBD produces unbiased estimates of *h*^2^ in relatively homogeneous samples. For phenotypes simulated from common CVs, unbiased estimates of 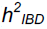 were also obtained using shorter cM thresholds. Results were similar for different relatedness thresholds (Figs S3 & S4) and for larger numbers of CVs (Fig. S5), although 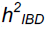 appeared to be biased upwards in phenotypes with 10,000 common CVs and long IBD length thresholds in the low stratification subsample (Fig. S5).

**Figure 1.**
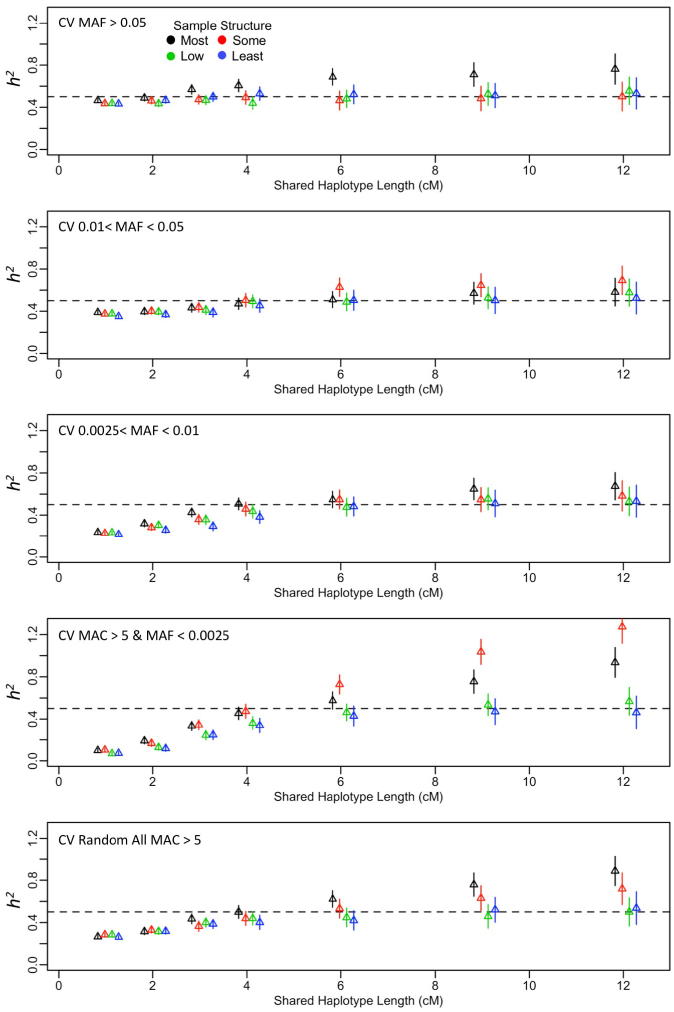
GREML-SC using an IBD-GRM. 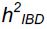 estimates (mean ± 95% CI from 400 replicates). X axis indicates the IBD shared haplotype length threshold for the IBD-GRM. Phenotypes with 1,000 CVs randomly drawn from the MAF range specified in each panel. Different colors indicate degree of stratification in the sample. Relatedness cutoff of 0.05 used.

In the two most stratified samples, we observed upward biases at long cM IBD thresholds, particularly for the rarest CVs (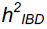> 1). This bias remained when using higher or lower relatedness thresholds (Figs. S3-4), and with 10,000 CVs (Fig S5).Controlling for 70 additional PCs or with additional PCs from very rare variants did not correct for the upward bias in very rare CV phenotypes, though inclusion of PCs did correct for bias in common CV phenotypes (Fig. S6). Furthermore, this bias was not mitigated by summing genetic length (cM) of IBD segments for calculating the GRM rather than physical length (Fig. S7) nor when using a two-GRM model, with one IBD-GRM calculated from even-numbered chromosomes and the second from odd-numbered chromosomes (Fig. S8-S9). Fitting a larger number of IBD-GRMs (e.g., one per chromosome) would better capture all the long-range correlations and might better mitigate the bias, but this approach is impractical for GREML-IBD in real data because the low variance of *D*_*ij*_ creates estimation problems. Thus, stratification has strong impacts on GREML-IBD estimates of heritability that we were unable to control for.

To explore why stratification had such strong influences on 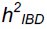 we first examined the correlations of off-diagonal GRM elements between the odd chromosome GRMs and even chromosome GRMs. Stratification clearly led to stronger long-range correlations, as did, in most subsamples, longer IBD thresholds for the GRM (Fig. S10). In the two least stratified subsamples, the correlation of even chromosome IBD-GRMs with odd chromosome WGS SNP-GRMs, estimated from either common or very rare WGS variants, was weak, and did not change drastically with increasing cM thresholds. There were stronger correlations overall in the two most stratified subsamples, especially between even chromosome IBD-GRMs and odd chromosome GRMs built from either IBD segments or from very rare WGS variants. Thus, stratification induced long-range correlations, such that *D*_*ij*_ for a pair of individuals at one chromosome predicted rare variant sharing at other chromosomes, which can presumably lead to over-estimation of 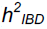 due to rare CVs being redundantly tagged by IBD sharing.

In simulations with odd chromosome CVs and IBD-GRMs calculated from even chromosomes only, we observed upward biases in 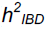 estimates for long IBD thresholds that were particularly severe in stratified samples with rare odd-chromosome CVs (Fig. S11). This pattern of results was similar to the pattern observed in our primary simulations (Fig. 1), consistent with the explanation that the upward biases in 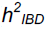 for rare CVs we observed at long IBD thresholds was due to long-range, redundant tagging of CVs in stratified samples. Note that the simulated *h*^2^ for the even chromosomes was 0. Because there is more recent common ancestry within than between subpopulations, there is more sharing of long IBD segments—and importantly more sharing of rare (recently arisen) causal variants. Consequently, due to stratification, long, shared IBD segments at one genomic location weakly predict not only sharing of long IBD segments, but also sharing of rare variants and shorter IBD segments, at other genomic locations. This redundant tagging of rare causal variants across the genome in stratified samples presumably leads to inflated 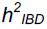 estimates. The same phenomenon has been described 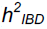 in the context of stratification (Yang *et al.*, 2011; Speed *et al.*, 2012), although the bias is less extreme and, because the variance of *A*_*ij*_ elements is much greater than the variance of *D*_*ij*_, is more easily alleviated by fitting multiple GRM models.

### Simulated Phenotypes – GREML-SNPs+IBD

The second model we tested was GREML-SNPs+IBD, which included a common SNP-GRM and the IBD-GRM. For phenotypes with 1,000 or 10,000 CVs, the total heritability 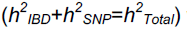 was unbiased in the two least stratified subsamples regardless of the CV MAF range (Fig. S12, S13). However, 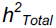 was again increasingly over-estimated in the two most stratified samples for very rare CV phenotypes. As expected, partitioning the variance to each of the GRMs, GREML-SNPs+IBD attributed more of the phenotypic variance to the common SNP-GRM when the CVs were common, and more of the variance to the IBD-GRM when the CVs were rarer (Figs. S14-S15). For common CV phenotypes, the variance attributable to the common SNP-GRM was overestimated by ~20%, which is consistent with previous findings for a common SNP-GRM based on the Axiom array positions and occurs because CVs in the common bin have higher average MAF than the SNPs on the Axiom array (Evans *et al.*, 2017). Interestingly, this overestimate was balanced by a negative variance estimate attributed to the IBD-GRM, such that the total estimated heritability was unbiased at ~0.5 (Figs. S12-S15). Nevertheless, 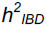 continued to be overestimated for very rare CV phenotypes in structured samples.

### Simulated Phenotypes – GREML-LDMS+IBD

Our third model included 16 imputed variant GRMs that were MAF- and LD-stratified, and the IBD-GRM. We found that across subsamples, GREML-LDMS+IBD produced unbiased 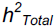 estimates with either 1,000 CVs or 10,000 CVs across all CV MAF ranges (Figs. S16-S17). Partitioning the variance among GRMs revealed that for the rare and very rare CV phenotypes, the IBD-GRM explained a small amount of the variance, but was near-zero otherwise (Figs. S18-S19).

When we excluded the rarest MAF bin from the model, leaving 12 imputed variant GRMs plus the IBD-GRM, GREML-LDMS+IBD produced unbiased 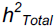 estimates with either 1,000 CVs or 10,000 CVs across all CV MAF ranges in subsamples with little or no stratification (Figs. 2, S20). However, with increased stratification, 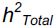 estimates were again overestimated for very rare CV phenotypes in the context of stratification. Partitioning the variance into that attributable to the LDMS imputed-variant GRMs and the IBD-GRM showed that, in unstratified samples, most of the genetic variance was attributable to the LDMS GRMs for CV MAF ranges > 0.0025 while the IBD-GRM captured the genetic variance for very rare CV MAF ranges (0.0003-0.0025) (Figs. 3, S21). While the variance attributed to the LDMS GRMs was never overestimated, that attributed to the IBD-GRMs at longer IBD thresholds was overestimated, resulting in total heritability estimates > 1 for the rarest CV phenotypes in the presence of stratification.

**Figure 2.**
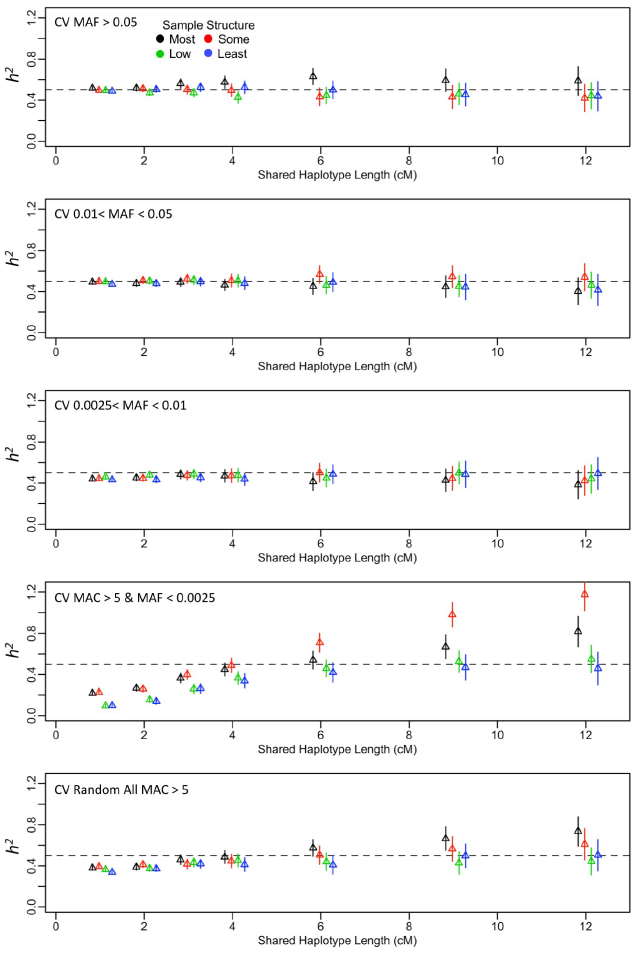
GREML-LDMS+IBD model. This model had 13 components, 12 LD & MAF-stratified GRMs using imputed genome-wide variants, and one GRM from IBD shared haplotypes. Total *h*^2^ estimates are shown (mean ± 95% CI from 400 replicates). X axis indicates the different IBD shared haplotype length thresholds for the IBD-GRM. Phenotypes with 1,000 CVs randomly drawn from the MAF range specified in each panel. Different colors indicate degree of stratification in the sample. Relatedness cutoff of 0.05 used.

**Figure 3.**
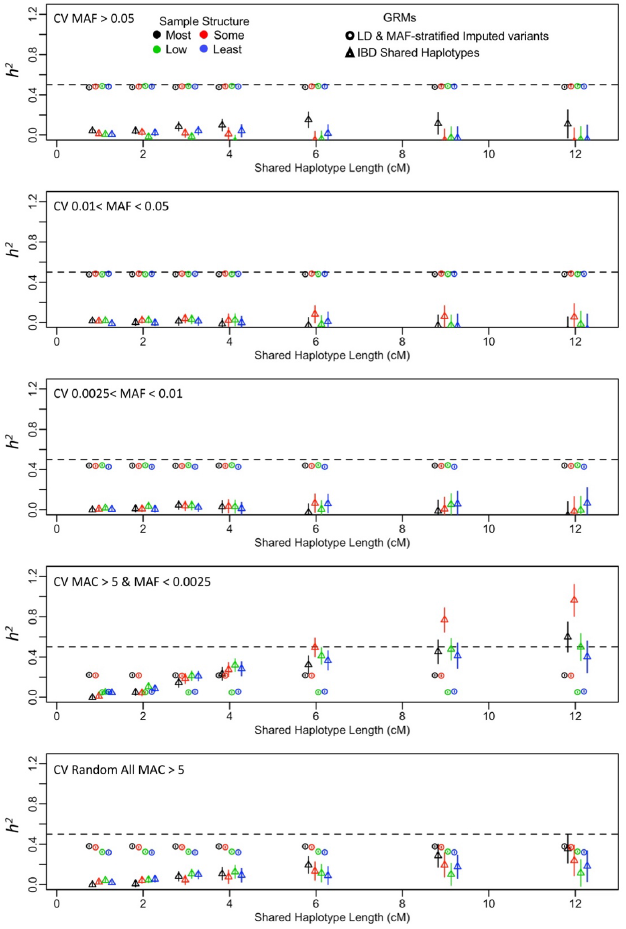
GREML-LDMS+IBD. This model had 13 components, 12 LD & MAF-stratified GRMs using imputed genome-wide variants, and one GRM from IBD shared haplotypes. Separate *h*^2^ estimates for each component are given by the symbols (mean± 95% CI from 400 replicates). Note that the “Imputed LDMS” symbol represents the sum of the imputed LDMS GRM variance estimates. X axis indicates the different IBD shared haplotype length thresholds for the IBD-GRM. Phenotypes with 1,000 CVs randomly drawn from the MAF range specified in each panel. Different colors indicate degree of stratification in the sample. Relatedness cutoff of 0.05 used.

### Real Phenotypes from the UK Biobank

Using GREML-IBD, 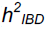 for height (but not for BMI) increased with longer minimum shared haplotype length, did not stabilize at longer segment thresholds, and appeared upwardly biased, similar to what we observed in stratified samples in our simulations (Fig. 4a, Table S1). The 95% CIs increased with longer minimum IBD length, as expected given the lower variance in *D*_*ij*_ at longer segment thresholds. For comparison, 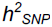 estimates from approaches using only SNPs are also presented in Table S1.

**Figure 4.**
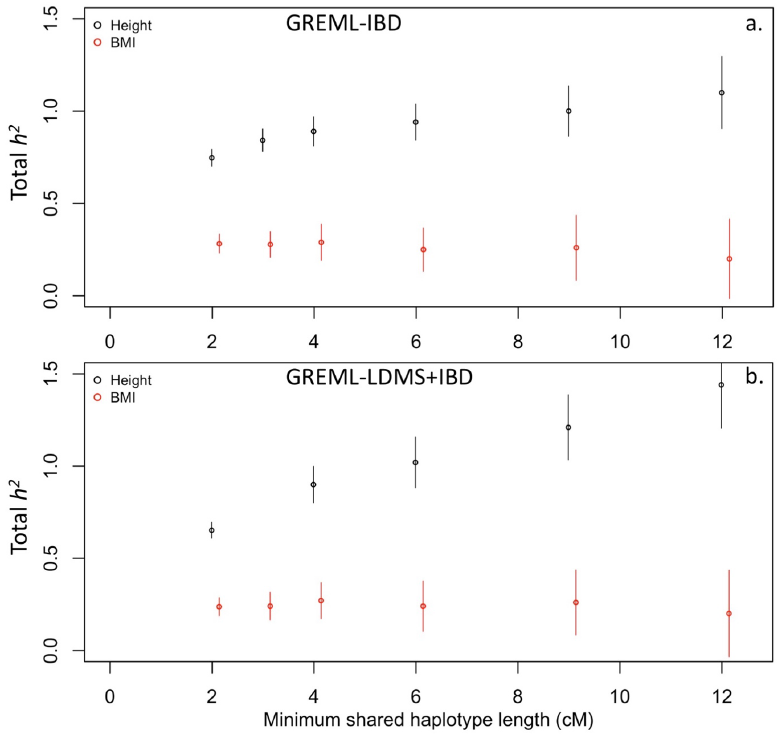
Total heritability estimates for three continuous traits in the UK Biobank. (a) GREML-IBD, which had a single IBD-GRM. (b) GREML-LDMS+IBD for three continuous traits in the UK Biobank. This model had 13 components, 12 LD & MAF-stratified GRMs using imputed genome-wide variants, and one GRM from IBD shared haplotypes. Total *h*^2^ estimates are shown (± 95% CI). X axis indicates the different IBD shared haplotype length thresholds for the IBD-GRM. Relatedness cutoff of 0.05 used.

Using either GREML-SNPs+IBD or GREML-LDMS+IBD, we found similar patterns of increasing 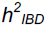 estimates with longer minimum IBD length for height, but the pattern was less extreme, and 95% CIs were generally smaller (Fig. 4b, Table S1).Results for GREML-LDMS+IBD either including the rarest MAF category or excluding it were similar: height 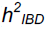 estimates increased from 0.75 to 1.1 across the range of minimum IBD lengths we examined. This increase in 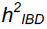 was due to increasing estimates of variance attributable to the IBD-GRM rather than to the imputed variant SNP-GRMs (Fig. S22, Table S1). BMI 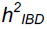 estimates were again ~0.2-0.3, though at longer minimum IBD length thresholds the standard errors were large, and the 95% CI overlapped 0 (Table S1).

Interestingly, inclusion of the IBD-GRM in addition to the SNP-GRM or LDMS-GRMs often improved model fit and resulted in a lower AICc (Table S1). Often the lowest AICc was found with shorter IBD minimum length thresholds. For instance, for height, the minimum AICc was found when using all LD- and MAF-stratified imputed variant GRMs and the IBD-GRM with a 3cM minimum IBD length threshold (Table S1), while AICc increased with longer length thresholds. Thus, while increasing the minimum length threshold led to unreasonable and uninterpretable total heritability estimates, at shorter IBD length thresholds, the inclusion of the IBD-GRM was preferred. This indicates that some additional variance remains to be explained over using only imputed-variant GREML-LDMS and that inclusion of the IBD-GRM led to models that better explained the observed phenotypic similarity, perhaps reflecting the effect of CVs that are not well captured by imputed variants.

## DISCUSSION

We present here the most thorough assessment to-date of an IBD-based heritability estimation approach. The interest in using IBD information in classically unrelated samples to estimate heritability arises from the potential to estimate the full narrow-sense heritability without the confounding of effects shared within families that can bias estimates when close relatives are used, and without the downward bias in estimation when CVs are rare or poorly tagged by SNPs. We demonstrated that GREML-IBD can produce unbiased heritability estimates in realistic whole-genome SNP data so long as there is little genetic stratification in the sample. Moreover, although we showed only marginal improvement, at best, over imputed variant-based approaches, IBD-based approaches should do increasingly better than ones based on imputed SNPs in estimating heritability if CVs are even rarer (MAF < .0003) than we could simulate here. That said, no estimation based on genomic sharing can capture variation due to CVs that occur only once in the sample.

While IBD-based approaches are appealing in principle, our study highlights two important drawbacks. First, stratification can bias heritability estimates upward, depending on the allele frequencies of CVs. The effect of stratification is strong when CVs are very rare, and is not controlled by inclusion of a large number of PC covariates, the typical approach to controlling such effects (Price *et al.*, 2010), or even PCs derived from very rare variants (Mathieson and McVean, 2012). Similar overestimates have been observed in a related method that used sharing at predefined, segregating haplotypes (Bhatia *et al.*, 2016). Overestimates appear to stem from redundant tagging by long IBD segments of very rare CVs as well as shorter IBD segments, particularly in stratified samples. Previous studies using IBD-based approaches (Zuk *et al.*, 2012; Browning and Browning, 2013) used isolated, homogeneous populations, which should mitigate this source of bias. Our simulation results suggest somewhat less homogenous samples, such as those of general northern/western European ancestry, can be used to derive unbiased heritability estimates so long as there are no additional confounding factors.

Second, the standard error (SE) of the 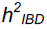 estimate is large due to the very low variance in IBD sharing among unrelated individuals in large, non-founder populations. For example, for height in the UK Biobank when using GREML-LDMS+IBD, total heritability SE≥0.053 for minimum IBD lengths ≥ 6cM, largely due to the IBD-GRM variance component SE. However, using just the imputed variant GREML-LDMS approach SE=0.015. Thus, while the GREML-LDMS+IBD may have accounted for more of the genetic variance, it did so with substantially lower precision. Very large sample sizes will be required to reach high levels of precision. Taken together, it is not clear whether the increased variance explained, arising from capturing rare CVs with IBD-based GRMs, outweighs the very large increase in standard errors and the increased potential for bias due to stratification or other factors we did not model here.

### Heritability of Real Complex Traits

Our results from real UK Biobank data for height demonstrate the potential for additional biases of an IBD-based approach that were not captured in our simulation. The estimates of total heritability for height increased with minimum IBD cM length, and were much greater than other reported estimates (e.g., Yang *et al.*, 2015; Evans *et al.*, 2017). This was unexpected given that the UK Biobank sample was similar to simulated data with respect to stratification. It is possible that the CVs, particularly rare CVs, underlying height are more geographically stratified than those that influence BMI. Indeed, evidence suggests that across Europe, genetic variance in height is more geographically structured than BMI, though both traits are more genetically structured than a neutral, drift-only model (Robinson *et al.*, 2015). However, environmental variation in BMI across Europe appears to be stronger than genetic differentiation (Robinson *et al.*, 2015). Thus, for height, neutral structure (shared long IBD segments) may covary with geographical variation in the CVs themselves, and lead to inflation of 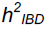. While consistent with our observed likely upward bias in height 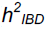 but not BMI 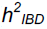, we cannot be certain that divergent selection is driving these patterns.

Vertically-transmitted non-genetic effects, shared common environmental effects, and assortative mating may also confound estimates of 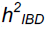. Estimates of 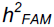 using close relatives can be altered by these factors (Eaves *et al.*, 1978; Martin *et al.*, 1978; Coventry and Keller, 2005;; Zuk *et al.*, 2012). It is currently unknown how GREML-based estimates, and IBD-based approaches in particular, are affected by assortative mating. Common environmental effects, which can induce similarity across highly extended pedigrees, would be confounded with IBD sharing, and are therefore a potential source of bias in IBD-based estimates. Such extended pedigree environmental similarity would be difficult to simulate using our simulation method, and so we did not explore how such an effect might bias 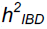. Our results in the UK Biobank data suggest that our assumption that removing close relatives (relatedness < 0.05) would mitigate shared environment confounding may require further investigation. The use of lower relatedness thresholds may alleviate the problem, but lower relatedness thresholds decrease the sample size and variance of IBD sharing and therefore further exacerbate the already high standard errors of these estimates. Rare variants are more differentially confounded by stratification than common variants, and typical approaches using PCA may not fully correct for such confounding (Mathieson and McVean, 2012). Extremely rare SNPs, as with long IBD segments, will co-segregate along extended pedigrees,and future work must focus on the role of confounding between familial and environmental effects and rare variants or long IBD segments.

While we cannot conclude with certainty which factors led to the apparent bias in height 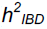, estimates of 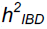 for BMI were more stable and also in line with previous reports. They suggest that BMI *h*^2^ is roughly 0.25-0.3, with up to 5% of the total phenotypic variance due to very rare or otherwise poorly-imputed variants that are captured by the IBD-GRM (see Table S1). As estimates from classical twin design studies range from 0.4-0.8, this suggests that much of the family-based estimates are due to shared environment, assortative mating, or non-additive genetic variance, supported by extended twin design variance estimates (Coventry and Keller, 2005; Keller and Coventry, 2005). This also suggests that little unexplained variance remains for BMI, as estimates of BMI 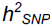 from recent studies range from 0.21 (Locke *et al.*, 2015) to 0.27 (Yang *et al.*, 2015).

Our findings may also offer context to the observed heritability estimates reported by several other studies that used haplotype-based approaches. Browning and Browning (2013) reported 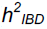 for BMI of 0, with standard error of 0.16 (height was not measured), although their upper 95% CI estimate is not inconsistent with a true *h*^2^ of 0.25-0.3. This low estimate may simply be due to sampling variance, arising from the small number of individuals (5,402) in the Finnish sample they used. Zaitlen et al. (2013) used IBD among close relatives to derive estimates of 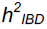 of 0.69 for height and 0.42 for BMI. As discussed by the authors, these estimates may be upwardly biased due to common environmental and non-additive genetic effects.

### Conclusions

Identical-by-descent haplotypes in common between a pair of chromosomes capture sharing at all variants that existed along their length in the last common ancestor. The ability to estimate such IBD segments using SNP data means that there is potential to estimate narrow-sense heritability of traits in a way that should be unbiased by factors that bias SNP or family-based estimates. We conclude that IBD-based estimates can be used to obtain estimates of the near full narrow-sense heritability. However, IBD-based estimates are imprecise and very sensitive to stratification. Moreover, when we estimated 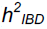 in real data, we observed unexpected biases that appeared similar to those that we had observed in more stratified samples in our simulation, which suggests that there are biases in real data that we were not able to adequately capture in our simulation. Taken together, these factors diminish the appeal of IBD-based approaches for estimating heritability, especially when compared to approaches that use imputed variants, such as GREML-LDMS. Nevertheless, until whole genome sequence data is feasible for the large sample sizes required for *h*^2^ estimation from genotype data, IBD-based estimates may be able to capture the rarest CVs better than imputation. In particular, though larger and more diverse reference panels are becoming available, isolated populations may not be well-represented. IBD-based approaches offer a method to capture rare genome-wide variants not represented in imputation reference panels, and these isolated, homogeneous populations may also be the most advantageous for IBD-based heritability estimation due to the larger variance in IBD sharing.

## ACKNOWLEDGEMENTS

This work utilized the Janus supercomputer, which is supported by the National Science Foundation (award number CNS-0821794), the University of Colorado Boulder, the University of Colorado Denver, and the National Center for Atmospheric Research, and is operated by the University of Colorado Boulder. We thank the participants of the individual HRC cohorts. This research has been conducted using the UK Biobank Resource. We thank the Keller and Vrieze lab groups, the Institute for Behavioral Genetics, and Sean Caron. This study was funded by NIH R01MH100141 (MCK), NHMRC grants 1078037 (PMV) and 1113400 (PMV and JY), and Sylvia & Charles Viertel Charitable Foundation Senior Medical Research Fellowship (JY).

## Haplotype Reference Consortium: http://www.haplotype-reference-consortium.org/

Gonçalo Abecasis, David Altshuler, Carl A Anderson, Andrea Angius, Jeffrey C Barrett, Sonja Berndt, Michael Boehnke, Dorrett Boomsma, Kari Branham, Gerome Breen, Chad M Brummett, Fabio Busonero, Harry Campbell, Peter Campbell, Andrew Chan, Sai Chen, Emily Chew, Massimiliano Cocca, Francis S Collins, Laura J Corbin, Francesco Cucca, Petr Danecek, Sayantan Das, Paul I W de Bakker, George Dedoussis, Annelot Dekker, Olivier Delaneau, Marcus Dorr, Richard Durbin, Aliki-Eleni Farmaki, Luigi Ferrucci, Lukas Forer, Ross M Fraser, Timothy Frayling, Christian Fuchsberger, Stacey Gabriel, Ilaria Gandin, Paolo Gasparini, Christopher E Gillies, Arthur Gilly, Leif Groop, Tabitha Harrison, Andrew Hattersley, Oddgeir L Holmen, Kristian Hveem, William Iacono, Amit Joshi, Hyun Min Kang, Hamed Khalili, Charles Kooperberg, Seppo Koskinen, Matthias Kretzler, Warren Kretzschmar, Alan Kwong, James C Lee, Shawn Levy, Yang Luo, Anubha Mahajan, Jonathan Marchini, Steven McCarroll, Mark I McCarthy, Shane McCarthy, Matt McGue, Melvin McInnis, Thomas Meitinger, David Melzer, Massimo Mezzavilla, Josine L Min, Karen L Mohlke, Richard M Myers, Matthias Nauck, Deborah Nickerson, Aarno Palotie, Carlos Pato, Michele Pato, Ulrike Peters, Nicola Pirastu, Wouter Van Rheenen, J Brent Richards, Samuli Ripatti, Cinzia Sala, Veikko Salomaa, Matthew G Sampson, David Schlessinger, Robert E Schoen, Sebastian Schoenherr, Laura J Scott, Kevin Sharp, Carlo Sidore, P Eline Slagboom, Kerrin Small, George Davey Smith, Nicole Soranzo, Timothy Spector, Dwight Stambolian, Anand Swaroop, Morris A Swertz, Alexander Teumer, Nicholas Timpson, Daniela Toniolo, Michela Traglia, Marcus Tuke, Jaakko Tuomilehto, Leonard H Van den Berg, Cornelia M van Duijn, Jan Veldink, John B Vincent, Uwe Volker, Scott Vrieze, Klaudia Walter, Cisca Wijmenga, Cristen Willer, James F Wilson, Andrew R Wood, Eleftheria Zeggini, He Zhang

## WHI Acknowledgment

The WHI program is funded by the National Heart, Lung, and Blood Institute, National Institutes of Health, U.S. Department of Health and Human Services through contracts HHSN268201600018C, HHSN268201600001C, HHSN268201600002C, HHSN268201600003C, and HHSN268201600004C. The authors thank the WHI investigators and staff for their dedication, and the study participants for making the program possible. A full listing of WHI investigators can be found at: http://www.whi.org/researchers/Documents%20%20Write%20a%20Paper/WHI%20Investigator%20Long%20List.pdf

## CONFLICT OF INTEREST

The authors declare no conflicts of interest.

## DATA ARCHIVING

Data are from the Haplotype Reference Consortium and the UK Biobank and can be accessed through those resources.

## REFERENCES

Bhatia G, Gusev A, Loh P-R, Finucane HK, Vilhjalmsson BJ, Ripke S, et al. (2016). Subtle stratification confounds estimates of heritability from rare variants. bioRxiv: 048181.

Bjelland DW, Lingala U, Patel P, Jones M, Keller MC (2017). A fast and accurate method for detection of IBD shared haplotypes in genome-wide SNP data. Eur J Hum Genet 25: 617–624.

Browning SR, Browning BL (2013). Identity-by-descent-based heritability analysis in the Northern Finland Birth Cohort. Hum Genet 132: 129–138.

Bulik-Sullivan BK, Loh P-R, Finucane HK, Ripke S, Yang J, Consortium SWG of the PG, et al. (2015). LD Score regression distinguishes confounding from polygenicity in genome-wide association studies. Nat Genet 47: 291–295.

Burnham KP, Anderson DR (2002). Model Selection and Multi-Model Inference, Second Edi. Springer New York.

Chang CC, Chow CC, Tellier LC, Vattikuti S, Purcell SM, Lee JJ, et al. (2015). Second-generation PLINK: rising to the challenge of larger and richer datasets. Gigascience 4: 7.

Coventry WL, Keller MC (2005). Estimating the Extent of Parameter Bias in the Classical Twin Design: A Comparison of Parameter Estimates From Extended Twin-Family and Classical Twin Designs. 8: 214–223.

Das S, Forer L, Schönherr S, Sidore C, Locke AE, Kwong A, et al. (2016). Next-generation genotype imputation service and methods. Nat Genet 48: 1284–1287.

Delaneau O, Zagury J-F, Marchini J (2013). Improved whole-chromosome phasing for disease and population genetic studies. Nat Methods 10: 5–6.

Eaves LJ, Last KA, Young PA, Martin NG (1978). Model-fitting approaches to the analysis of human behaviour. Heredity 41: 249–320.

Evans LM, Tahmasbi R, Vrieze SI, Abecasis GR, Das S, Bjelland DW, et al. (2017). Comparison of methods that use whole genome data to estimate the heritability and genetic architecture of complex traits. bioRxiv: 115527.

Falconer DS, Mackay TFC (1996). Introduction to Quantitative Genetics, Longman Limited: Harlow, Essex, England.

Gusev A, Lowe JK, Stoffel M, Daly MJ, Altshuler D, Breslow JL, et al. (2009). Whole population, genome-wide mapping of hidden relatedness. Genome Res 19: 318– 326.

Hayes BJ, Visscher PM, Goddard ME (2009). Increased accuracy of artificial selection by using the realized relationship matrix. Genet Res (Camb) 91: 47–60.

Keller MC, Coventry WL (2005). Quantifying and addressing parameter indeterminacy in the classical twin design. Twin Res Hum Genet 8: 201–213.

Lee SH, DeCandia TR, Ripke S, Yang J, Sullivan PF, Goddard ME, et al. (2012). Estimating the proportion of variation in susceptibility to schizophrenia captured by common SNPs. Nat Genet 44: 247–250.

Locke AE, Kahali B, Berndt SI, Justice AE, Pers TH, Day FR, et al. (2015). Genetic studies of body mass index yield new insights for obesity biology. Nature 518: 197– 206.

Lynch M, Walsh B (1998). Genetics and Analysis of Quantitative Traits. Sinauer Associates: Sunderland, MA.

Martin NG, Eaves LJ, Kearsey MJ, Davies P (1978). The power of the classical twin study. Heredity 40: 97–116.

Mathieson I, McVean G (2012). Differential confounding of rare and common variants in spatially structured populations. Nat Genet 44: 243–6.

McCarthy S, Das S, Kretzschmar W, Delaneau O, Wood AR, Teumer A, et al. (2016). A reference panel of 64,976 haplotypes for genotype imputation. Nat Genet 48: 1279–1283.

Polderman TJC, Benyamin B, de Leeuw CA, Sullivan PF, van Bochoven A, Visscher PM, et al. (2015). Meta-analysis of the heritability of human traits based on fifty years of twin studies. Nat Genet 47: 702–709.

Price AL, Helgason A, Thorleifsson G, McCarroll SA, Kong A, Stefansson K (2011). Single-tissue and cross-tissue heritability of gene expression via identity-by-descent in related or unrelated individuals. PLoS Genet 7.

Price AL, Zaitlen N a, Reich D, Patterson N (2010). New approaches to population stratification in genome-wide association studies. Nat Rev Genet 11: 459–63.

Ripke S, Neale BM, Corvin A, Walters JTR, Farh K-H, Holmans PA, et al. (2014). Biological insights from 108 schizophrenia-associated genetic loci. Nature 511: 421–427.

Robinson MR, Hemani G, Medina-Gomez C, Mezzavilla M, Esko T, Shakhbazov K, et al. (2015). Population genetic differentiation of height and body mass index across Europe. Nat Genet 47: 1357–1362.

Speed D, Hemani G, Johnson MR, Balding DJ (2012). Improved heritability estimation from genome-wide SNPs. Am J Hum Genet 91: 1011–1021.

Sudlow C, Gallacher J, Allen N, Beral V, Burton P, Danesh J, et al. (2015). UK Biobank: An Open Access Resource for Identifying the Causes of a Wide Range of Complex Diseases of Middle and Old Age. PLoS Med 12: 1–10.

Tenesa A, Haley CS (2013). The heritability of human disease: estimation, uses and abuses. Nat Rev Genet 14: 139–149.

Visscher PM, Brown MA, McCarthy MI, Yang J (2012). Five years of GWAS discovery. Am J Hum Genet 90: 7–24.

Visscher PM, Hill WG, Wray NR (2008). Heritability in the genomics era-concepts and misconceptions. Nat Genet 9: 255–266.

Visscher PM, Medland SE, Ferreira M a R, Morley KI, Zhu G, Cornes BK, et al. (2006). Assumption-free estimation of heritability from genome-wide identity-by-descent sharing between full siblings. PLoS Genet 2: e41.

Wakeley J (2009). Coalescent Theory: An Introduction. Roberts and Company: Greenwood Village, CO.

Wood AR, Esko T, Yang J, Vedantam S, Pers TH, Gustafsson S, et al. (2014). Defining the role of common variation in the genomic and biological architecture of adult human height. Nat Genet 46: 1173–86.

Yang J, Bakshi A, Zhu Z, Hemani G, Vinkhuyzen AAE, Lee SH, et al. (2015). Genetic variance estimation with imputed variants finds negligible missing heritability for human height and body mass index. Nat Genet 47: 1114–20.

Yang J, Benyamin B, McEvoy BP, Gordon S, Henders AK, Nyholt DR, et al. (2010). Common SNPs explain a large proportion of the heritability for human height. Nat Genet 42: 565–569.

Yang J, Lee SH, Goddard ME, Visscher PM (2011). GCTA: A tool for genome-wide complex trait analysis. Am J Hum Genet 88: 76–82.

Yang J, Manolio TA, Pasquale LR, Boerwinkle E, Caporaso N, Cunningham JM, et al. (2011). Genome partitioning of genetic variation for complex traits using common SNPs. Nat Genet 43: 519–25.

Zaitlen N, Kraft P, Patterson N, Pasaniuc B, Bhatia G, Pollack S, et al. (2013). Using extended genealogy to estimate components of heritability for 23 quantitative and dichotomous traits. PLoS Genet 9.

Zuk O, Hechter E, Sunyaev SR, Lander ES (2012). The mystery of missing heritability: Genetic interactions create phantom heritability. Proc Natl Acad Sci U S A 109: 1193–8.

